# Developmental plasticity of hermaphrodite sperm production across environments in *Caenorhabditis elegans*

**DOI:** 10.1101/2025.06.18.660282

**Authors:** Clotilde Gimond, Nausicaa Poullet, Anne Vielle, Emilie Demoinet, Christian Braendle

## Abstract

Many organisms show flexible resource allocation to adjust for optimal reproductive investment across different environments. How such reproductive plasticity occurs in hermaphroditic organisms—allocating resources to both oocytes and sperm—are central questions of sex allocation research. Self-fertilizing hermaphrodites of the androdioecious nematode *Caenorhabditis elegans* exhibit a sequential transition from spermatogenesis to oogenesis, so the extent of self-sperm production determines both fertilization onset and lifetime reproductive potential under selfing. Despite this key role, it remains largely unclear whether such sequential hermaphrodites flexibly adjust sperm production to optimize self-fertilization across different environments. Here we directly quantified plasticity in *C. elegans* hermaphrodite self-sperm production in diverse experimental environments. We found that: (a) Sperm production was developmentally plastic, but such changes did not consistently translate into changes in self-progeny number, suggesting *C. elegans* self-fecundity is often oocyte-limited rather than sperm-limited; (b) Contrary to expectations, plastically increased sperm production did not delay the onset of fertilization across various environments; (c) Subtle environmental challenges, such as mild dietary restriction, did not affect sperm production but had a significant impact on developmental time, age at reproductive maturity, and germline proliferation. This emphasizes the relative environmental insensitivity of sperm production compared to other reproductive traits in hermaphrodites. (d) Plasticity in sperm and germline traits varied by genetic background, with notable differences between the laboratory strain N2 and wild strains. These findings contribute to our understanding of reproductive plasticity in *C. elegans* and the developmental plasticity of sex allocation in sequential hermaphrodites.

## INTRODUCTION

Phenotypic plasticity of developmental, morphological and physiological traits shapes an organism’s reproductive life history depending on the environmental context. Such reproductive plasticity encompasses plasticity in sex allocation, which is predicted to reflect the optimal investment of resource to female and male functions. One central question is how reproductive plasticity occurs in hermaphroditic organisms, where the same individual must coordinate the allocation of limited resources to both female (oocytes) and male (sperm) function. In general, sex allocation theory on hermaphrodites predicts that resource allocation to male function depends on the degree of selfing (self-fertilization), with expected low investment to male function under high selfing and hermaphrodites producing the minimal amount of self-sperm to fertilize all oocytes (Charnov 1982). Theoretical and empirical studies on plasticity in hermaphrodite sex allocation have been conducted across diverse taxa, but have predominantly focused on simultaneous hermaphrodites and their sex allocation plasticity in response to environmental variation—particularly changes in social context, such as shifts in local mate competition (Charlesworth and Charlesworth 1981; Locher and Baur 2002; Schärer and Ladurner 2003; Schärer 2009; Singh and Schärer 2022). In contrast, little is known about the environmental sensitivity of sex allocation in sequential protandrous hermaphrodites that initially produce self-sperm and then irreversibly switch to oocyte production, allowing for self-fertilization.

Here we focus on developmental plasticity of sex allocation in selfing hermaphrodites of the androdioecious (male-hermaphrodite) nematode *Caenorhabditis elegans*. Androdioecy in this nematode with males (XO) and hermaphrodites (XX) is evolutionarily derived from a male-female breeding system (dioecy) (Kiontke *et al*. 2011). Hermaphrodites are altered females, with their male function restricted to producing self-sperm, so they cannot mate and exchange sperm with one another (Hirsh *et al*. 1976; Byerly *et al*. 1976; Ward and Carrel 1979; Kimble and Ward 1988). Cross-fertilization with males occurs, yet rarely, with self-fertilization being the predominant reproductive mode of *C. elegans* in natural populations (Barriere and Felix 2005; Cutter *et al*. 2019). During *C. elegans* larval development, hermaphrodite germ cells in the two symmetrical gonad arms initially differentiate into sperm, followed by a switch to oogenesis (Fig. 1A). This type of sequential hermaphroditism affects two critical, interconnected determinants of reproductive fitness. First, the sperm-oocyte switch is irreversible, so that the number of self-sperm produced will pre-determine and limit the potential for self-fertilization. *C. elegans* fecundity is thus considered to be limited by the number of available self-sperm rather than oocytes. This contrasts a majority of organisms, in which oocytes, as the much larger, more costly gamete type, represent the limiting factor (Bateman 1948; Charnov 1982). Second, due to the sequential nature of the sperm-oocyte switch, the extent of self-sperm production will critically determine the onset of fertilization, with increased self-sperm production leading to a delay in reproductive maturity (Hodgkin and Barnes 1991; Barker 1992; Cutter 2004; Chasnov 2011; Braendle and Paaby 2024). The sperm-oocyte switch thus underpins a fundamental trade-off between generation time and lifetime fecundity under selfing (Fig. 1B).

**Figure 1.**
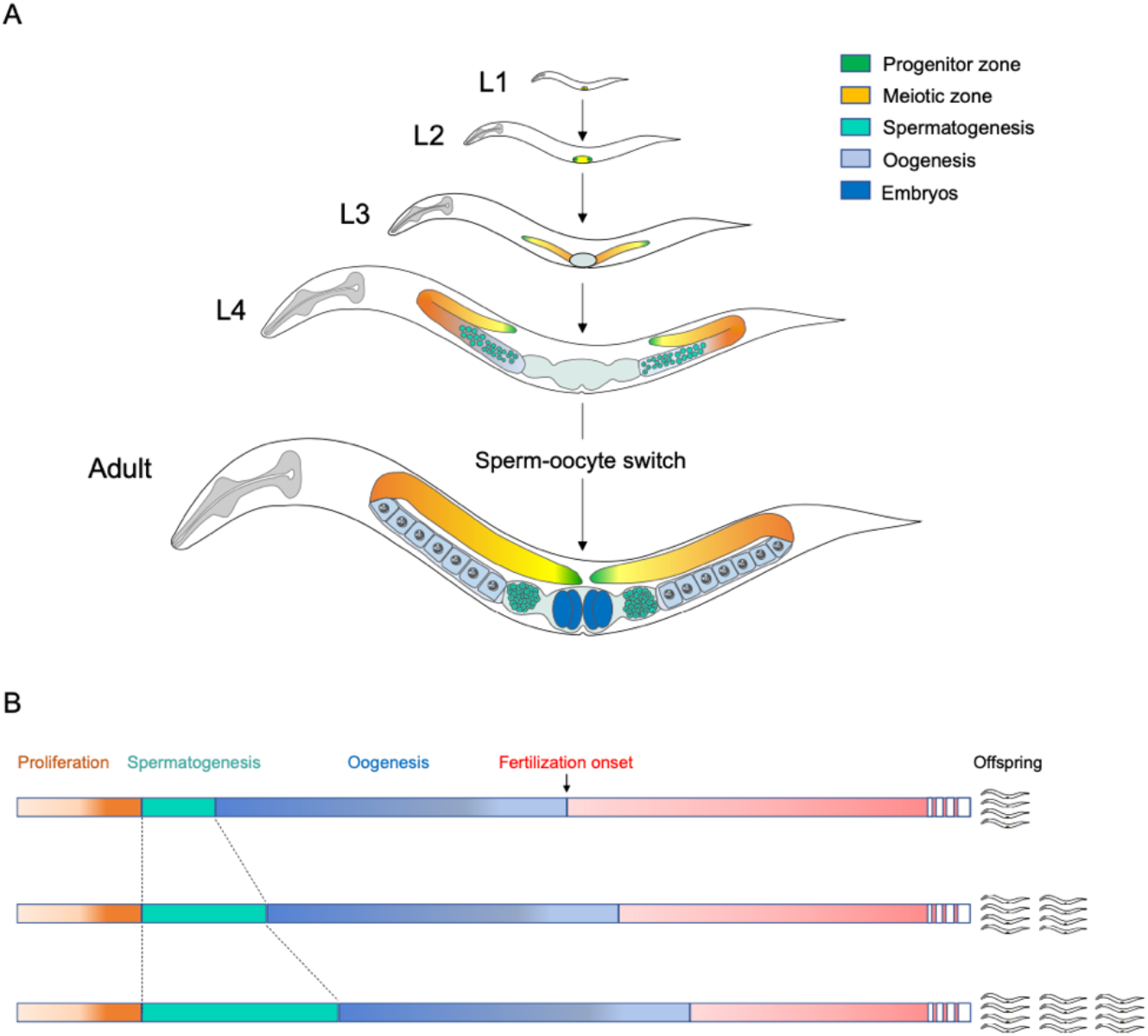
Sequential production of sperm and oocytes in the *C. elegans* hermaphrodite. (A) Larval germline development of the *C. elegans* hermaphrodite through the four larval stages (L1 to L4) and adult stage (Kimble and Crittenden 2007). The distal proliferative region of the germline, also termed progenitor zone (PZ), contains mitotic germ cells representing germ stem cell pool (Hubbard and Schedl 2019). Germ cell proliferation begins in the L1 stage. Cells exiting the PZ enter the transition zone (green-yellow to orange) to meiosis, first differentiating as sperm (L3-L4), then as oocytes. The adult *C. elegans* germline comprises two symmetrical U-shaped gonad arms connected to a central uterus. The distal region of each arm contains mitotically dividing germ cells, including stem cells, maintained by signalling from the Distal Tip Cell (DTC, green). Mature oocytes are fertilized upon passing through the spermatheca, and resulting embryos develop within the uterus. (B) The sperm-oocyte switch underlies a fundamental trade-off between generation time and lifetime fecundity under selfing. The sperm-oocyte switch reflects a fundamental trade-off between generation time and lifetime fecundity under selfing. Prolonged spermatogenesis delays the switch, increasing generation time but potentially enhancing total fecundity. Conversely, an earlier switch shortens generation time at the possible cost of reduced selfing capacity.

As shown by theoretical models evaluating this specific trade-off – using growth rate as the relevant fitness measure for *C. elegans* – the predicted optimum of hermaphrodite self-sperm production is close to what has been observed in experiments and is thus consistent with sex allocation theory (Barker 1992; Cutter 2004). Under favourable laboratory conditions with *ad libitum* food, a self-fertilizing *C. elegans* hermaphrodite produces a maximum of ~250-300 self-sperm, which corresponds closely to the number of lifetime self-progeny, indicating that self-fertilization is very efficient (Ward and Carrel 1979; Hodgkin and Barnes 1991; Singson 2001; Cutter 2004). A hermaphrodite provided with sperm by mating with males can produce more than 1000 offspring, hence, fecundity of exclusively self-fertilizing *C. elegans* hermaphrodites is sperm-limited in such laboratory settings (Ward and Carrel 1979; Hodgkin and Barnes 1991; Cutter 2004).

Despite its central role in hermaphrodite life history, little is known about the environmental sensitivity of self-sperm production and its link to plasticity in selfing capacity across diverse conditions. Although, over a century ago, Emile Maupas had already reported that *C. elegans* reared under “bad conditions” produced only 30–40 spermatids, compared to up to 240 under optimal conditions (Maupas 1900), only few studies have since directly assessed environmental effects on hermaphrodite sperm production, e.g. in response to temperature (Poullet *et al*. 2015), bacterial food source (Mishra *et al*. 2023), male pheromone (Faerberg *et al*. 2024) or after passage through the dauer stage (Ow *et al*. 2018). To gain a more comprehensive view of sperm plasticity, we directly quantified hermaphrodite self-sperm production across a range of environmental conditions varying in nutrient availability, microbial food source, and abiotic stressors—possible factors reflective of the natural, highly variable and complex, habitats of *C. elegans* (Félix and Braendle 2010). Our primary objective was to assess the degree of plasticity in self-sperm production and evaluate how this plasticity corresponds to reproductive output under exclusive selfing. This allowed us to determine whether self-fecundity is sperm-limited across different environmental conditions. We then examined three specific factors—nutrient limitation, dauer passage, and microbial food source—that influence self-sperm number, to explore how plasticity in sperm production relates to changes in timing and dynamics of germline development and lifetime reproductive output. In addition to assessing environmental influences, we examined how genetic background modulates environmental effects on sperm production, allowing us to compare the differences in reproductive plasticity between laboratory-adapted and wild strains.

## RESULTS

### Plasticity of hermaphrodite sperm production in different environments

We quantified hermaphrodite sperm production in the laboratory reference strain N2 and the genetically divergent wild strain CB4856 (Hawaii) across ten environmental conditions that varied in microbial food source, temperature, and chemical composition of the growth medium. While sperm production varied significantly among environments, the extent of plasticity was similar between the two strains (Fig. 2A). Lifetime offspring production (Fig. 2B), measured in parallel from animals of the same experimental cohort, generally correlated with sperm number across environments, with higher sperm production translating into higher offspring output (Fig. 2C, D). Growth in liquid culture markedly reduced lifetime offspring numbers in both strains without affecting sperm production. Additionally, in CB4856, ethanol or osmotic stress led to a stronger decoupling of sperm and offspring production compared to N2 (Fig. 2C, D). In general, plastic changes in hermaphrodite sperm production tended to mirror shifts in self-offspring output. Yet, in certain stressful environments, sperm numbers substantially exceeded offspring counts, pointing to limitations in oocyte production or ovulation. Still, we cannot exclude that some environments may impair sperm function, as previously reported for high temperature (Petrella 2014; Poullet *et al*. 2015).

**Figure 2.**
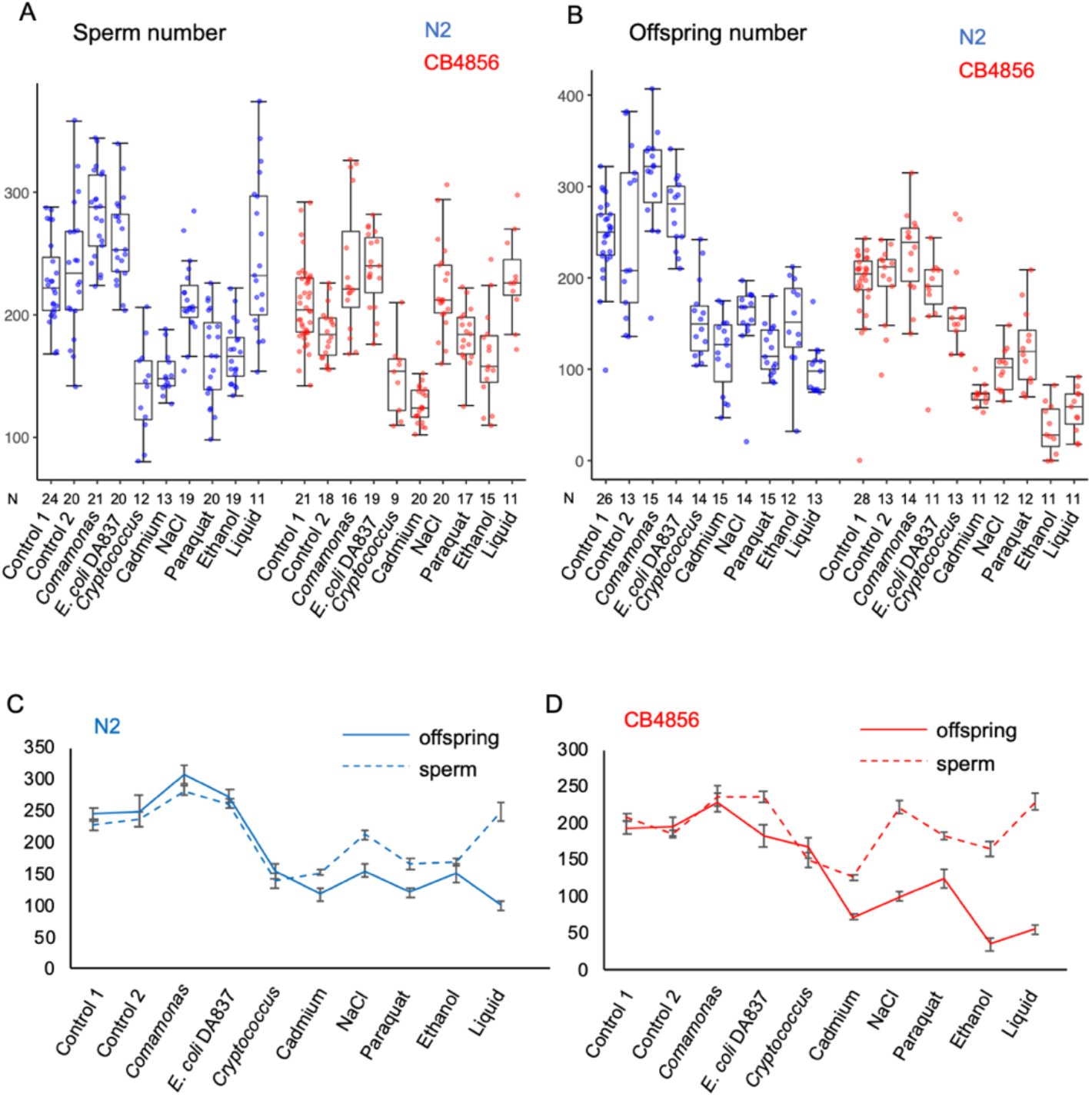
Plasticity of hermaphrodite sperm production across ten environments. Number of sperm (A) and (B) number of offspring produced in N2 and CB4856 grown in various environments. See Materials and Methods for environments and their corresponding controls (two-way ANOVA for sperm, genotype: F_1,354_ = 16.61, P < 0.0001; environment: F_9,354_ = 34.12, P < 0.0001; genotype x environment: F_9,354_ = 3.37, P = 0.0006; N = 9-41; two-way ANOVA for offspring, genotype: F_1,267_ = 89.16, P < 0.0001; environment: F_9,267_ = 54.85, P < 0.0001; genotype x environment: F_9,267_ = 6.05, P < 0.0001; N = 11,28). (C, D) Corresponding reaction norms for the same data.

### Effects of nutrient deprivation on hermaphrodite sperm production

We first tested how several mutants (*eat-2, rsks-1, pept-1*), causing severe physiological starvation despite the presence of food impact sperm and offspring production (Lakowski and Hekimi 1998; Meissner *et al*. 2004; Korta *et al*. 2012; Grimbert *et al*. 2018). These three mutants, all displaying extreme delays in larval development and onset of reproductive maturity, showed strongly reduced sperm and offspring numbers (Fig. 3A). In all cases, sperm numbers substantially exceeded offspring output, supporting the notion that hermaphrodite reproduction under severe nutrient limitation is constrained by oocyte production rather than sperm availability (Goranson *et al*. 2005; Braendle and Paaby 2024). Evidence that oogenesis becomes strongly reduced in these mutants comes from the markedly reduced number of cells observed in the distal progenitor zone (Fig. 3B)

**Figure 3.**
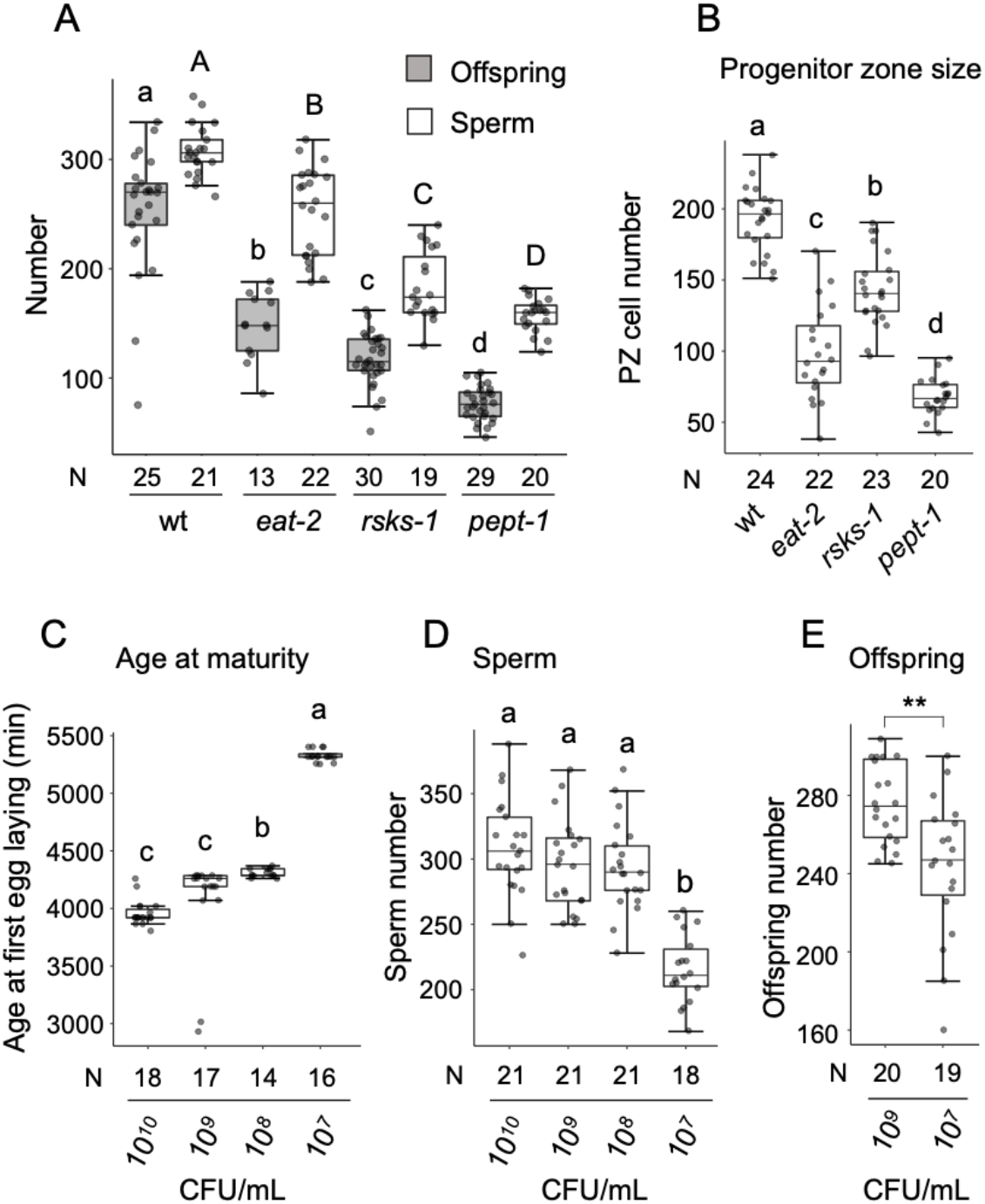
Effects of nutrient deprivation on hermaphrodite sperm production. (A) Sperm and offspring in DR mutants (ANOVA, F_3,93_ = 119.09, P < 0.0001, N=13-30). (B) Number of germ cells in the distal progenitor zone (PZ) of mutants (ANOVA, F_3,83_ = 106.00, P < 0.0001, N=20-24). (C) Age at maturity varied significantly among bacterial dilution treatments (ANOVA, F_3,61_ = 77.30, P < 0.0001, N=14-18). Time is indicated in minutes from the egg-laying window. (D) Sperm number varied significantly among bacterial dilution treatments (ANOVA, F_3,77_ = 32.83, P < 0.0001, N=18-21). (E) Lifetime offspring number differed significantly between two of the bacterial dilution treatments (ANOVA, F_1,37_ = 11.82, P = 0.0015, N=19-20). Bars labelled with different letters represent statistically significant differences (Tukey’s HSD test); asterisks indicate significance levels: P < 0.05 (*), P < 0.01 (), P < 0.001 (*).

We next investigated how dietary restriction (DR) affects hermaphrodite sperm production in the N2 strain by exposing animals to four concentrations of *E. coli* OP50. As anticipated, lower food availability delayed overall development, thereby postponing reproductive maturity (Fig. 3C). DR treatment also significantly reduced self-sperm numbers, but only at the lowest food concentration (10^7^ CFU/mL) (Fig. 3D), which was accompanied by a corresponding decline in offspring production (Fig. 3E).

To assess genetic variation in the plasticity of sperm production under reduced nutrient availability, we compared the N2 strain to three wild strains using the strongest DR treatment (107 CFU/mL). This treatment markedly delayed development across all four strains (data not shown) and consistently reduced the number of nuclei in the distal progenitor zone of the wild strains without affecting sperm numbers (Fig. 4A, B). In contrast, the N2 laboratory strain showed no reduction in cells of the distal progenitor zone but exhibited a significant decrease in sperm number under nutrient deprivation (Fig. 4A, B). Hence, the impact of nutrient deprivation on sperm production and reproductive output in *C. elegans* is strongly genotype-dependent, with strain-specific differences such as the pronounced fecundity reduction observed in CB4856. We confirmed that dietary restriction frequently reduces oogenic germline proliferation and growth (Korta *et al*. 2012; Ow *et al*. 2018; Hubbard and Schedl 2019; Baugh and Hu 2020). However, this reduction does not necessarily extend to sperm production. Our results suggest that hermaphrodite sperm production is often unaffected by nutrient deprivation (Fig. 3D and 4B), unless the restriction is particularly severe, as observed in mutants inducing very extreme nutrient deprivation (Fig. 3A).

**Figure 4.**
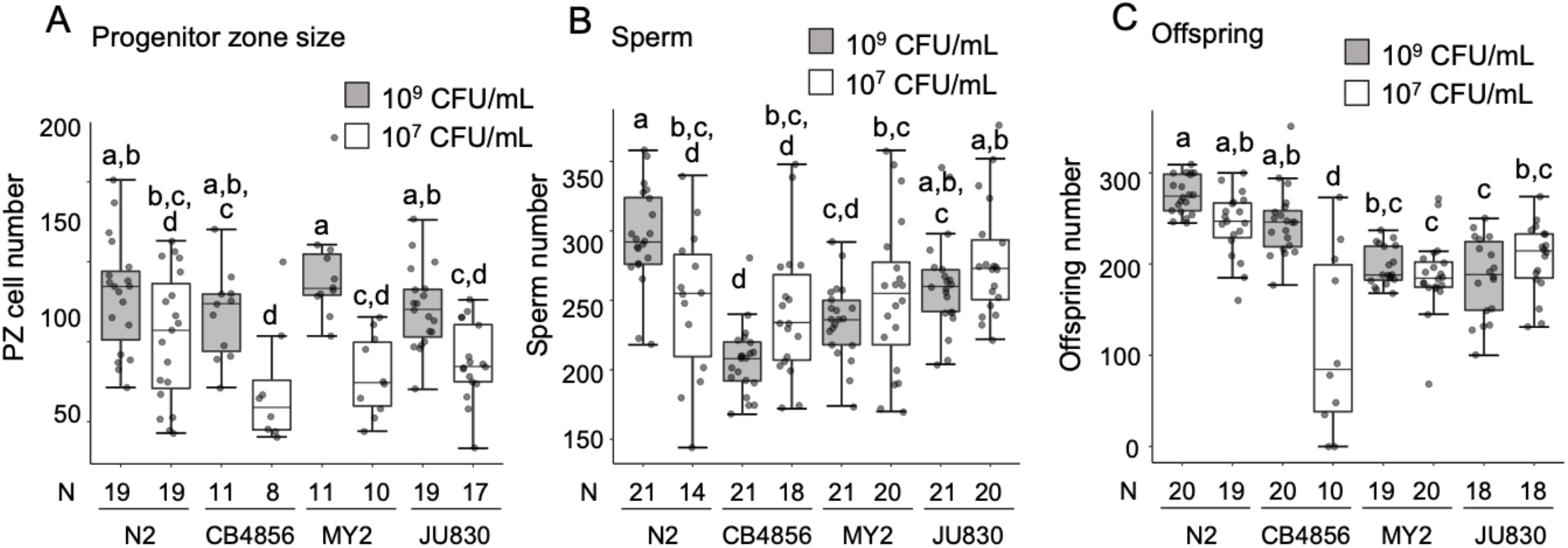
Genetic variation in the plasticity of sperm production under nutrient deprivation. (A) Number of germ cells in the distal progenitor zone (PZ) (two-way ANOVA, genotype: F_3,106_ = 2.27, P = 0.0084; environment: F_1,106_ = 49.15, P < 0.0001; genotype x environment: F_3,106_ = 1.80, P = 0.1512; N=8-19). (B) Sperm number (two-way ANOVA, genotype: F_3,148_ = 12.09, P < 0.0001; environment: F_1,148_ = 0.59, P = 0.4446; genotype x environment: F_3,148_ = 6.91, P = 0.0002; N=14-21). (C) Lifetime offspring number (two-way ANOVA, genotype: F_3,136_ = 20.83, P < 0.0001; environment: F_1,136_ = 29.41, P < 0.0001; genotype x environment: F_3,136_ = 20.17, P < 0.0001; N=10-20). Bars labelled with different letters represent statistically significant differences (Tukey’s HSD test).

### Effects of dauer passage on hermaphrodite sperm production

*C. elegans* dauer induction is a central life history decision governed by multiple environmental cues, including ascaroside pheromone concentration (a proxy for population density), food availability and quality, or temperature (Baugh and Hu 2020; Braendle and Paaby 2024). Theoretical models predict that dauer passage, linked to dispersal and colonization of new food patches, should favour increased self-sperm production to optimize reproductive success (Chasnov 2011). However, empirical support for this prediction is mixed: postdauer adults can exhibit either increased or decreased self-offspring, depending on the specific dauer-inducing cue and duration of the dauer stage (Kim and Paik 2008; Hall *et al*. 2010, 2013; Ow *et al*. 2018, 2021; Webster *et al*. 2018). Most of these studies did not directly quantify sperm numbers, leaving the effects of dauer passage on self-sperm production unresolved.

We therefore examined how different dauer-inducing cues and time spent in dauer influence sperm production in postdauer hermaphrodites. In line with previous findings, N2 postdauer adults produced more self-offspring (Fig. 5A) and correspondingly more sperm (Fig. 5B) when dauer was induced via high population density (growth on egg-white plates). This increase was especially pronounced in individuals that remained in dauer for only one day (Fig. 5B). In contrast, the wild strain CB4856 showed a different response: sperm production was unchanged after 1 day in dauer but decreased following 5 days (Fig. 5B). Regardless of dauer duration, CB4856 postdauer adults consistently exhibited lower self-offspring numbers than non-dauer controls (Fig. 5A). While dauer passage had variable effects on sperm production, it consistently reduced oogenic germline proliferation in both strains under all conditions tested, as evidenced by a decrease in cells within the distal progenitor zone (Fig. 5C).

**Figure 5.**
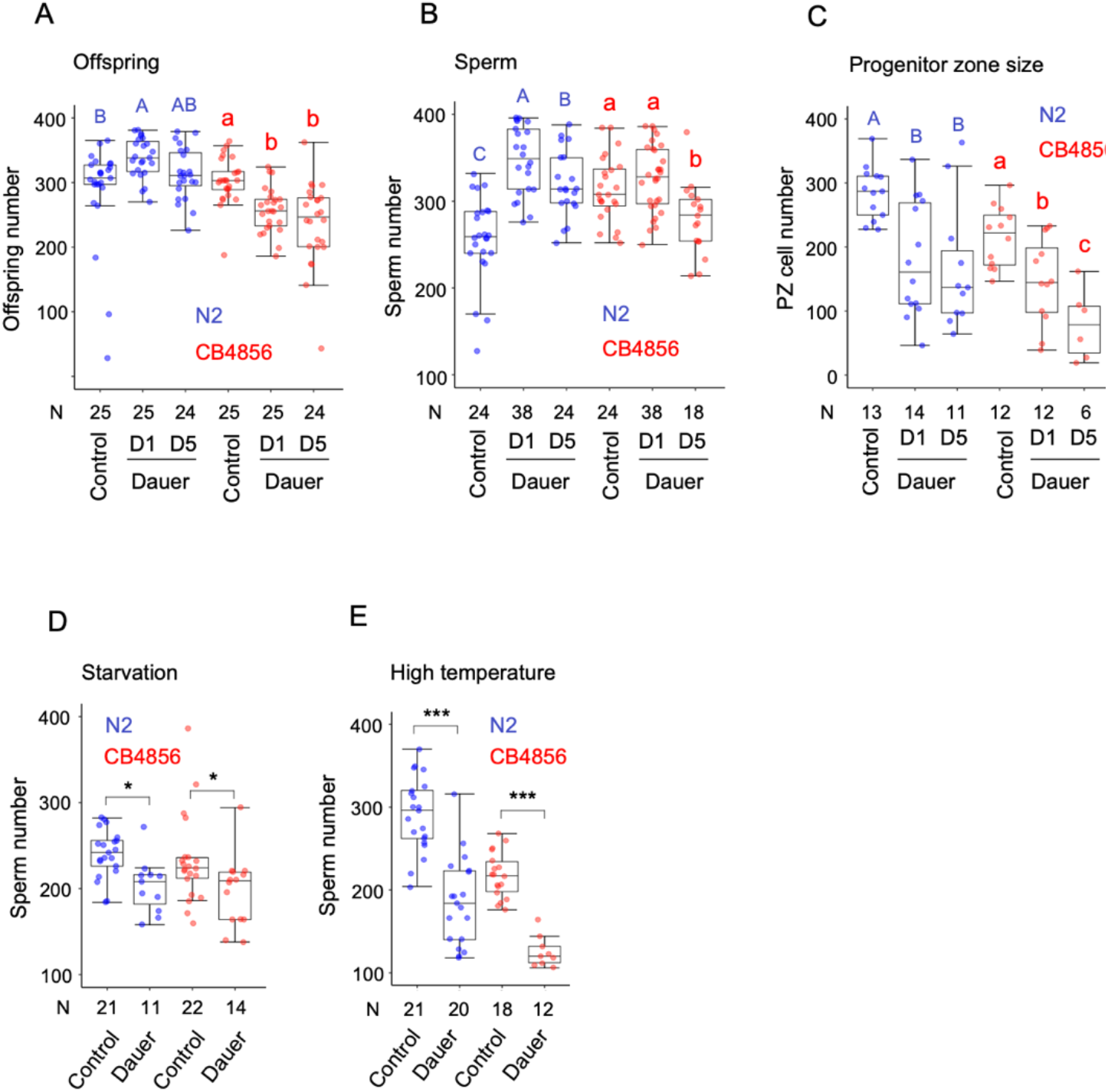
Effects of dauer passage on hermaphrodite sperm production. (A-C) Effects of dauer passage induced by high population density (egg-white plates) in strains N2 and CB4856. Values with different letters indicate statistically significant differences (P < 0.05). (A) Lifetime offspring number; N2: ANOVA, F_2,71_ = 5.29, P = 0.0072, N = 24–25; CB4856: ANOVA, F_2,71_ = 10.84, P < 0.0001, N = 24–25. (B) Sperm counts; N2: ANOVA, F_2,83_ = 42.67, P < 0.0001, N = 24–38; CB4856: ANOVA, F_2,77_ = 10.18, P = 0.0001, N = 18– 38. (C) Number of germ cells in the distal progenitor zone (PZ); N2: ANOVA, F_2,35_ = 8.33, P = 0.0011, N = 11–14; CB4856: ANOVA, F_2,27_ = 11.55, P = 0.0002, N = 6–12. (D) Effects on hermaphrodite sperm production following dauer passage induced by starvation (N2: Kruskal-Wallis Test, 𝒳^2^ = 8.98, P = 0.003; CB4856: Kruskal- Wallis Test, 𝒳^2^ = 6.67, P = 0.01). (E) Effects on hermaphrodite sperm production following dauer passage induced by high temperature (N2: Kruskal-Wallis Test, 𝒳^2^ = 22.55, P < 0.0001; CB4856: Kruskal- Wallis Test, 𝒳^2^ = 20.91, P < 0.0001). Asterisks indicate significant differences (*P < 0.05, **: P < 0.001, ***: P < 0.0001).

To further investigate the influence of environmental cues, we tested whether alternative dauer-inducing conditions (starvation and elevated temperature) also impact sperm production. Both treatments significantly reduced sperm numbers in postdauer individuals across both strains (Fig. 5D, E). Together, these findings demonstrate that sperm production following dauer passage is influenced by at least three interacting factors: genetic background, time spent in dauer, and the specific dauer-inducing cue.

### Effects of *Serratia marcescens* on hermaphrodite sperm production

Compared to the standard food sources *E. coli* OP50, several alternative sources significantly enhanced *C. elegans* sperm and offspring production, including *Comamonas* sp. DA1877 and *E. coli* DA837 (Fig. 2A, B). We observed similar positive effects with *Serratia marcescens*, a species that co-occurs with *C. elegans* in natural environments (Samuel *et al*. 2016) and is known for its pathogenicity under laboratory conditions (Mallo *et al*. 2002; Kurz *et al*. 2003). Intrigued by these seemingly positive fitness effects of a pathogenic food source, we quantified how *S. marcescens* affects reproduction by rearing animals exclusively on pure cultures of the (pathogenic) laboratory strain of *S. marcescens* Db10 (Mallo *et al*. 2002). Growth on *S. marcescens* significantly increased lifetime offspring number of *C. elegans* N2 and CB4856 relative to animals feeding on the standard diet, *E. coli* OP50 (Fig. 6A). This increase in reproductive output was coupled to a corresponding increase in sperm production (Fig. 6B) as well as accelerated development so that individuals feeding on *S. marcescens* reached reproductive maturity 2-3h earlier (Fig. 6C). Additionally, young adult hermaphrodites fed with *S. marcescens* exhibited a significant expansion of adult germ cell progenitor pools (Fig. 6D,E). Consistent with an earlier onset of reproductive maturity (Fig. 6C), *S. marcescens* accelerated germline progression and differentiation, as evidenced by multiple indicators—including the presence of spermatocytes, spermatids, anti-RME-2 staining, and/or mature oocytes— observed at distinct developmental stages (Fig. 6F). For instance, at the L4/adult moult, 20% of individuals fed with *S. marcescens* were positive for the early oocyte marker RME-2, compared to only 3% of those fed *E. coli OP50* (Fig. 6F). These results show that *C. elegans* developmental growth and reproduction are generally enhanced when feeding on *S. marcescens*. In particular, the *S. marcescens* Db10 strain improved all measured fitness-related traits in both N2 and CB4856 strains, suggesting it functions more as a beneficial nutritional source than a pathogen—despite its known harmful effects on post-reproductive lifespan. This apparent paradox is supported by previous findings indicating that *C. elegans* aversion to *S. marcescens* may reflect stage-specific susceptibility and early larvae are resistant to infection (Kurz *et al*. 2003), while later exposure impairs survival and reproduction (Zhang *et al*. 2005; Pradel *et al*. 2007).

**Figure 6.**
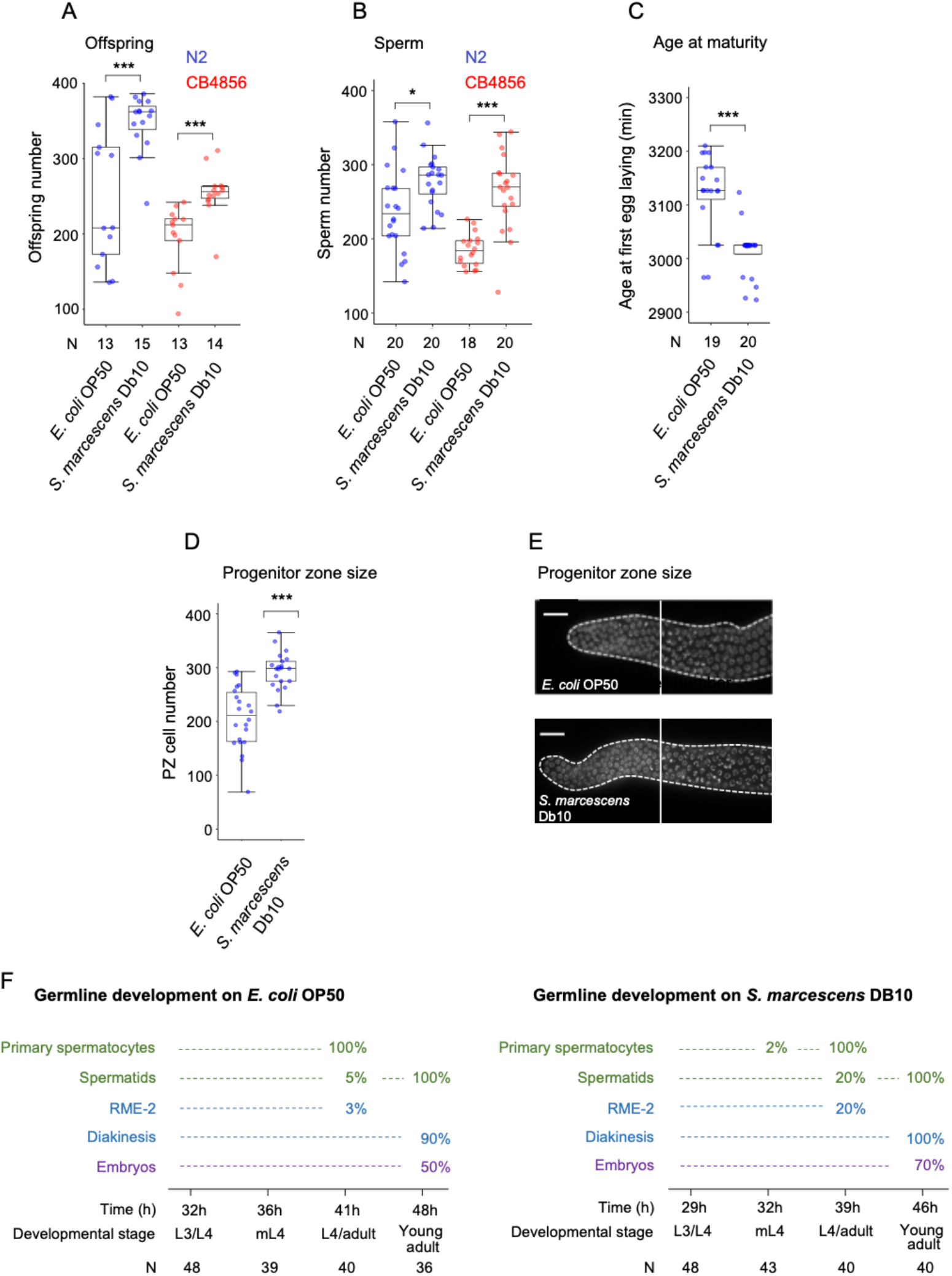
Effects of *Serratia marcescens* on hermaphrodite sperm production. (A) Effects of *Serratia marcescens* Db10 on lifetime offspring production in strains N2 and CB4856 (N2: Kruskal- Wallis Test, 𝒳^2^ = 7.91, P = 0.0049; CB4856: Kruskal- Wallis Test, 𝒳^2^ = 15.08, P < 0.0001). (B) Effects of *Serratia marcescens* Db10 on sperm production in strains N2 and CB4856 (N2: Kruskal-Wallis Test, 𝒳^2^ = 6.54, P = 0.0105; CB4856: Kruskal- Wallis Test, 𝒳^2^ = 19.50, P < 0.0001). (C) Effects of *Serratia marcescens* Db10 on reproductive timing (strain N2): age at maturity (i.e. time from hatching to first egg laid) was reached significantly faster on *S. marcescens* compared to *E. coli*. Kruskal-Wallis Test, 𝒳^2^ = 16.79, P < 0.0001. (D) Number of germ cells in the distal progenitor zone (PZ) in young adult hermaphrodites (L4+24h) (ANOVA, F_1,41_ = 29.40, P < 0.0001). (E) DAPI-stained images of dissected gonad arms from young adults, showing the distal germline region. Dotted lines outline the germline; solid lines mark the boundary between the mitotic and meiotic zones (transition zone). (F) Quantification of germline progression and gamete maturation at defined developmental stages, normalized to absolute developmental time from L1 hatch. Transition nuclei, pachytene-stage cells, and spermatocytes were scored from DAPI-stained gonads. Oocyte progression was analyzed using RME-2 staining to detect early oocytes and DAPI nuclear morphology to identify mature oocytes at diakinesis (Poullet *et al*. 2015, 2016). For experimental details, see Materials and Methods section. Asterisks indicate statistically significant differences (*P < 0.05, **: P < 0.001, ***: P < 0.0001).

## DISCUSSION

Our results provide insight into how *C. elegans* hermaphrodites modulate sex allocation across different environments. While developmental plasticity of the germline in response to the environment, including plasticity in proliferation, apoptosis, oocyte maturation, and ovulation, is well documented (Narbonne and Roy 2006; Korta and Hubbard 2010; Hubbard *et al*. 2013; Sowa *et al*. 2015; Hubbard and Schedl 2019; Baugh and Hu 2020; Ow *et al*. 2021; Fausett *et al*. 2021; Bollen *et al*. 2023; Braendle and Paaby 2024; Kloock and Hubbard 2025), plasticity of self-sperm production has typically been only indirectly inferred by measuring offspring number. Here, by directly quantifying sperm numbers in self-fertilizing *C. elegans* hermaphrodites, we show that sperm production is indeed plastic. However, plastic changes in sperm production did not consistently predict changes self-fecundity, supporting the idea that *C. elegans* reproduction is often oocyte-limited, especially in suboptimal conditions (Goranson *et al*. 2005; Murray and Cutter 2011). Furthermore, we observed no consistent trade-off between sperm number and the timing of reproductive maturity. Both sperm production and broader reproductive plasticity showed strong genotype dependence.

### Hermaphrodite sperm production is plastic across environments

Severe environmental stressors, such as strong dietary restriction, cadmium, ethanol, oxidative stress, and exposure to *Cryptococcus* fungi, consistently reduced fecundity and were accompanied by decreased self-sperm production. In contrast, environments like liquid culture reduced offspring numbers without affecting self-sperm levels. Mild nutrient limitation significantly delayed development and reduced germline proliferation but did not alter sperm production. This suggests that sperm production is relatively insensitive to moderate stress, potentially due to its lower energetic cost compared to oocyte production. Such resilience may help maintain a baseline level of self-sperm to ensure reproductive assurance via selfing.

### No universally visible trade-off between sperm number and reproductive timing

As sequential hermaphrodites, *C. elegans* initiate spermatogenesis during larval development and switch to oogenesis upon entering adulthood. This temporal separation has led to the hypothesis that increasing self-sperm production delays the onset of reproduction. While such trade-offs have been revealed under specific genetic (e.g., germline sex determination mutants) or environmental conditions (e.g., alternative bacterial diets) (Hodgkin and Barnes 1991; Mishra *et al*. 2023; Braendle and Paaby 2024), our findings suggest this is not a universal pattern. For instance, in the N2 strain, strong nutrient deprivation delayed reproductive maturity and reduced sperm production. In contrast, exposure to *Serratia marcescens* Db10 accelerated development while increasing both sperm production and fecundity. These results indicate that resource allocation between male and female functions does not always constrain reproductive timing (Poullet *et al*. 2016). Yet, the absence of a visible trade-off does not necessarily mean it is absent. In some cases—particularly under suboptimal conditions—other physiological effects of environmental stress may obscure or override the expected trade-off, making it difficult to detect.

### Genetic variation in plasticity of hermaphrodite sperm production

Environmental effects on hermaphrodite sperm production differed between different wild-type strains, indicative of heritable variation in developmental plasticity of sperm production. Wild strains exhibited idiosyncratic responses in reproductive plasticity across environments, but N2 often behaved distinctly. Specifically, some environmental effects appeared strain-specific: while dauer passage increased sperm and offspring production in N2 (Hall *et al*. 2010, 2013; Ow *et al*. 2018, 2021), this response was absent in CB4856, which instead showed reductions in both traits under the same high-population treatment. Nevertheless, both strains exhibited decreased sperm counts following dauer induced by starvation or heat, aligning with previous findings (Ow *et al*. 2018).

## Conclusions

Our findings connect self-sperm plasticity to broader patterns of germline and life history plasticity in *C. elegans*, providing novel insights into sex allocation in a protandrous hermaphroditic species with sequential sperm-oocyte development. In particular, our study highlights hermaphrodite sperm production as a central, yet previously underappreciated, aspect of reproductive plasticity in *C. elegans*. While sperm production appears less environmentally sensitive than other germline traits (proliferation, as measured by the mitotic progenitor zone size), it is still subject to plastic modulation in response to severe or specific environmental factors. These data underscore the importance of resource availability and variable environmental conditions in sex allocation plasticity in *C. elegans*. The pronounced genotype-dependence in sperm and reproductive responses underscores the evolutionary divergence of reproductive strategies in natural *C. elegans* populations. It will be valuable to investigate whether consistent differences in hermaphrodite sperm production are found across distinct habitats or geographic regions. For example, do *C. elegans* strains or haplotypes that have spread globally and are commonly associated with human-modified environments and abundant food resources (Lee *et al*. 2019) show consistently elevated sperm production and greater selfing capacity? These questions remain unanswered. Addressing them will require detailed comparisons with newly sampled isolates from highly divergent, likely ancestral populations, particularly those from the Pacific region such as Hawaii (Crombie *et al*. 2019, 2022; Lee *et al*. 2021).

## MATERIALS AND METHODS

### Strains and culture conditions

For experimental procedures and materials, we used standard protocols of C. elegans research (Stiernagle 2006). The *C. elegans* strain N2 (UK) and the wild strains CB4856 (Hawaii), MY2 and JU830 (Germany) were kindly provided by Marie-Anne Félix. Mutant strains *eat-2(ad465)* (strain DA365), *rsks-1(ok1255)* (strain RB1206) and *pept-1(lg601)* (strain BR2742) were obtained from the *Caenorhabditis* Genetics Center (CGC). Animals were maintained at 20°C on 2.5% agar Nematode Growth Medium (NGM) plates seeded with the *E. coli* strain OP50 following standard procedures (Stiernagle 2006).

Animals for a given experiment were derived from the same maternal and grandmaternal environmental conditions without undergoing starvation. Unless indicated otherwise, experimental populations were age-synchronized by hypochlorite treatment and L1 arrest in M9 buffer (Stiernagle 2006). Resulting L1 individuals were randomly allocated to the different experimental treatments as detailed below. Animals examined in each experiment were grown using the same materials (NGM stock solutions, bacterial solutions, etc.) and different strains and phenotypes were always scored in parallel.

### Phenotyping protocols

#### Sperm production

Sperm numbers were counted in young adult hermaphrodites, i.e. adults with a maximum of 1 or 2 embryos in the uterus, as described previously (Poullet *et al*. 2015; Vielle *et al*. 2016; Gimond *et al*. 2019; Fausett *et al*. 2023). Briefly, animals were collected in M9 buffer (2.2 mM KH2PO4, 4.2mM Na2HPO4, 85 mM NaCl, 1 mM MgSO4) and fixed in cold methanol for 30 minutes at -20C, washed twice with PBSTw (PBS: phosphate-buffered saline, 137 mM NaCl, 2.7 mM KCl, 10 mM Na2HPO4, 2 mM KH2PO4, pH 7.4 containing 0.1% Tween20) and mounted on glass slides with Vectashield mounting medium supplemented with 4,6-diamidino-2-phenylindole (DAPI; Vector Labs). Images were taken at 60X magnification as Z-stacks covering the entire thickness of the gonad using an Olympus BX61 microscope with a CoolSnap HQ2 camera. We counted sperm number in one gonadal arm of a given individual by identifying DAPI-stained sperm nuclei in different focal planes through the specimen using ImageJ plugin Cell Counter, and doubled this number to derive an estimate of the total number of self-sperm produced by each hermaphrodite. When primary spermatocytes were still present, they were counted as four spermatids.

#### Offspring production

Lifetime self-offspring production (fecundity) was quantified across different environments and mutant backgrounds, following established protocols (Poullet *et al*. 2015). In parallel to animals selected for sperm number counts, additional hermaphrodites were isolated on individual plates of the corresponding environmental condition. Animals were then transferred daily to fresh plates until egg production ceased. The total number of offspring produced by each individual was obtained after adding counts obtained from each plate.

#### Germ cell nuclei number in the progenitor zone

Germ cell nuclei in the distal proliferative region, also termed progenitor zone (PZ), were counted as described previously using whole-body DAPI staining (Poullet *et al*. 2015, 2016).

#### RME-2 antibody staining

Oogenesis onset (Fig. 6F) was scored with RME-2 antibody staining of the germline (Grant and Hirsh 1999) following the same protocol as in a previous study (Poullet *et al*. 2016).

### Plasticity of hermaphrodite sperm production in different environments (Fig. 2 and Fig. 6)

#### Control conditions

We employed two control conditions. First, standard NGM plates (2.5% agar) seeded with an *E. coli* OP50 lawn were used as the primary control, named “Control 1” in Figure 2 (Stiernagle 2006). Second, to match the conditions used for testing other microbial strains, we included NGM plates without bactopeptone, indicated as “Control 2” in Figure 2. This modification limits bacterial overgrowth, which can interfere with the observation of nematodes, and serves as an appropriate baseline for assays involving microbial diets other than *E. coli* OP50.

#### Microbial food sources

In addition to the standard food source *E. coli* OP50, the following alternative microbial strains were used: *E. coli* DA837 (derived from *E. coli* OP50 but is more difficult for worms to eat) (Shtonda 2006), *Comamonas sp*. DA1877 (a bacterium on which *C. elegans* grows particularly well) (Avery 2003; MacNeil *et al*. 2013) as well as the pathogenic *Serratia marcescens* Db10 (Mallo *et al*. 2002; Kurz *et al*. 2003). In addition, we included a yeast strain *Cryptococcus kuetzingii* (ATCC42276), which can serve as a food source and affects diverse life history traits in *C. elegans* (Mylonakis *et al*. 2002).

Bacterial strains were cultured overnight on Lysogeny Broth (LB) agar plates at 37°C. A single colony was then used to inoculate LB liquid medium, which was incubated at 37°C with shaking. The yeast *Cryptococcus kuetzingii* was cultured on Yeast Extract Peptone Dextrose (YPD) plates and in YPD liquid medium at 30°C. To control microbial concentration, bacterial and yeast cultures were washed twice in S-basal medium (5.85 g/L NaCl, 1 g/L K2HPO4, 6 g/L KH2PO4, 5 mg/L cholesterol), and the optical density (OD) was measured at 600 nm. To prepare experimental plates, NGM plates lacking bactopeptone were used to limit microbial overgrowth, which can obscure nematode visibility. Plates were poured several days in advance and inoculated with 100 µL of microbial culture at 10^9^ CFU /mL. After inoculation, plates were incubated at room temperature for two days to allow for consistent bacterial lawn formation, then transferred to 4°C for storage (up to one month) until use in experiments.

#### NaCl (Osmotic stress)

We used a 5-fold increased NaCl concentration (250mM) compared to control NGM plates. The salt was directly added to the NGM medium before autoclaving (Salinas *et al*. 2006).

#### Paraquat (Oxidative stress)

Oxidative stress caused by the herbicide paraquat induces strong germ cell apoptosis (Salinas *et al*. 2006). A 1M stock solution was prepared, filtered and added to the NGM medium after autoclaving, for a final concentration of 0.5mM (Salinas *et al*. 2006; Fausett *et al*. 2021).

#### Cadmium

Cadmium induces strong germ cell apoptosis and reduces germ cell proliferation in a dose and time-dependant manner (Wang *et al*. 2008). A 100mM stock solution of cadmium was prepared, filtered and added to the NGM medium after autoclaving, for a final concentration of 25μM.

#### Ethanol

Ethanol plates were made according to a previous study (Davis *et al*. 2008), which showed that chronic ethanol exposure delays development and reduces both fecundity and lifespan. To prepare the plates, 95% ethanol was evenly pipetted onto OP50-seeded agar plates to achieve a final ethanol concentration of 0.25 M in the agar. Plates were then sealed with Parafilm and allowed to equilibrate at room temperature for at least 2 hours before transferring animals.

#### Liquid

1mL of S-basal (Stiernagle 2006) was added to regular NGM plates seeded with *E. coli* OP50. Plates were then sealed with Parafilm to avoid evaporation.

### Effects of nutrient deprivation on hermaphrodite sperm production (Fig. 3 and 4)

To evaluate how nutrient deprivation affects sperm production (Fig. 3), we tested a range of *E. coli* OP50 concentrations from 10^10^ to 10^7^ CFU/mL. The aim was to identify a dietary restriction (DR) condition that significantly impaired fitness without completely abolishing reproductive capacity. Based on these preliminary assessments, we selected 107 CFU/mL as the treatment and 10^9^ CFU/mL as the control condition. All traits across the four tested strains (Fig. 4) were measured in parallel within a single experimental setup. Animals were cultured on NGM plates lacking bactopeptone and seeded with either control (10^9^ CFU/mL) or treatment (10^7^ CFU/mL) *E. coli* OP50

### Effects of dauer passage on hermaphrodite sperm production (Fig. 5)

#### Dauer formation in response to high population density

Dauer formation was induced using high-density populations on egg-white supplemented plates at 25°C, based on a protocol adapted from D.L. Baillie and R.E. Rosenbluth (personal communication) (Hall *et al*. 2010; Ow *et al*. 2018), Briefly, one chicken egg white was added to 50 mL of boiling distilled water, homogenized in a blender for one minute, and 1 mL of the mixture was layered onto 55 mm NGM plates seeded with *E. coli* OP50. Plates were allowed to dry overnight. Approximately 5,000 hypochlorite-synchronized L1 larvae were transferred onto these plates and incubated at 25°C. After three days, the majority of animals on egg-white plates had entered dauer. In parallel, L3 larvae that had not entered dauer were picked the day after plating and transferred to fresh OP50-seeded NGM plates at 20°C as controls. To assess the effect of dauer duration, dauer larvae were collected from egg-white plates after 1, 5, or 12 additional days and isolated using 1% SDS treatment for 20 minutes. Following SDS selection, dauers were transferred to standard NGM plates seeded with *E. coli* OP50 and maintained at 20°C until they reached adulthood.

#### Dauer formation in response to starvation

The protocol for starvation-induced dauer formation followed a previously published protocol (Ailion and Thomas 2000). Experimental populations were established by placing two adult hermaphrodites on each plate and incubating them at 20°C. Four days after the bacterial lawn was completely depleted, 1% SDS was applied to the plate, and dauer larvae were scored after 20 minutes of exposure. Control animals (adults and L4 larvae) were collected from the plates prior to the onset of starvation. ***Dauer formation in response to high temperature:*** The protocol for temperature-induced dauer formation was based on previously published protocol (Ailion and Thomas 2000). Approximately 30 adults were allowed to lay eggs for 5 hours at room temperature. Plates were then shifted to 27°C to induce partial dauer formation. As a control, L3-stage larvae were picked after one day at 27°C and transferred to fresh seeded plates at 20°C. Dauer larvae were picked 44 hours after the egg-laying window and placed on standard NGM plates at 20°C for recovery to adulthood.

## Supporting information

Experimentaldatatables

## ACKNOWLEDGEMENTS

We thank the laboratories of Jonathan Ewbank, Marie-Anne Félix, Judith Kimble, Eleftherios Mylonakis for sharing nematode and microbial strains with us.

